# Landmark Kernel tICA for Conformational Dynamics

**DOI:** 10.1101/123752

**Authors:** Matthew P. Harrigan, Vijay S. Pande

## Abstract

Molecular dynamics simulations of biomolecules produce a very high dimensional time-series dataset. Performing analysis necessarily involves projection onto a lower dimensional space. *A priori* selection of projection coordinates requires (perhaps unavailable) prior information or intuition about the system. At best, such a projection can only confirm the intuition. At worst, a poor projection can obscure new features of the system absent from the intuition. Previous statistical methods such a time-structure based independent component analysis (tICA) and Markov state modeling (MSMs) have offered relatively unbiased means of projecting conformations onto coordinates or state labels, respectively. These analyses are underpinned by the propagator formalism and the assumption that slow dynamics are biologically interesting. Although arising from the same mathematics, tICA and MSMs have different strengths and weaknesses. We introduce a unifying method which we term “landmark kernel tICA” (lktICA) which uses a variant of the Nyström kernel approximation to permit approximate non-linear solutions to the tICA problem. We show that lktICA is equivalent to MSMs with “soft” states. We demonstrate the advantages of this united method by finding improved projections of (a) a 1D potential surface (b) a peptide folding trajectory and (c) an ion channel conformational change.

## 1 Introduction

Protein dynamics are responsible for carrying out the functions of life. Molecular dynamics (MD) is a powerful tool to understand dynamics on an atomistic scale by modeling and simulating each atom as a classical particle. Thanks to increasing computer power, simulations have modeled larger systems at longer timescales. Among other challenges, drawing interpretable conclusions from these increasingly high-dimensional and increasingly lengthy time-series data sets has been called in to focus ^1^. Markov State Models (MSMs) ^2–4^ and time-structure based independent component analysis (tICA) ^5–7^ have been introduced to address this challenge. These models are backed by a useful formalism: the transfer operator ^3^. This has given rise to a variational approach ^8,9^ which can be used to select the best models (hyperparameters) for a given dataset ^10^. Furthermore, it has been shown that tICA and MSMs solve the same problem in this formalism, namely numerical estimation of the transfer operator. The difference between the two is in choice of basis set ^11^. In particular, MSMs construct indicator-function basis functions over microstates, and tICA uses linear basis functions in the input coordinates. It is noteworthy that tICA was introduced primarily as an intermediate processing step for MSM construction, and only sometimes connected by this formalism ^5^.

Figure 1 shows an example potential energy function V(x) yielding four wells and a large barrier between the leftmost and rightmost wells. For a simple, low-dimensional potential energy surface, we can analytically calculate the equilibrium distribution *μ*(*x*) = exp[–*βν*(*x*)]/*Ζ*, where *Z* is the partition function Σexp[–*βν*(*x*)]. For a high-dimensional potential energy surface where quadrature integration is impossible we must use a different approach. For example, we can use molecular dynamics to sample conformations of a protein in solvent subject to a force field. A natural way of estimating the distribution 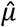 ≈ *μ* is by constructing a histogram (Figure 1a). Specifically, we partition the data into bins and count the number of data points in each bin. In addition to thermodynamic properties, we may also be interested in kinetic properties. We expect Brownian dynamics on this potential to yield three slow processes corresponding to transitions among the four wells. The slowest process is transfer of flux from the left two wells to the right two wells (Figure 1b). The transfer operator (and related propagator) formalism sets up a framework for estimating kinetics from data. The MSM is the kinetic analog of the thermodynamic histogram: we partition the data into discrete states and count transitions between states. Both the histogram (thermodynamics) and MSM (kinetics) describe smooth data with jagged bins, making estimation highly sensitive to shot noise, poor statistics, and binning protocol. To overcome this limitation, one might consider using a “smooth” estimator. One smooth analogue to the histogram is the kernel density estimator (KDE). In a similar vein, we seek a more sophisticated technique for smoothly estimating kinetics.

**Figure 1:**
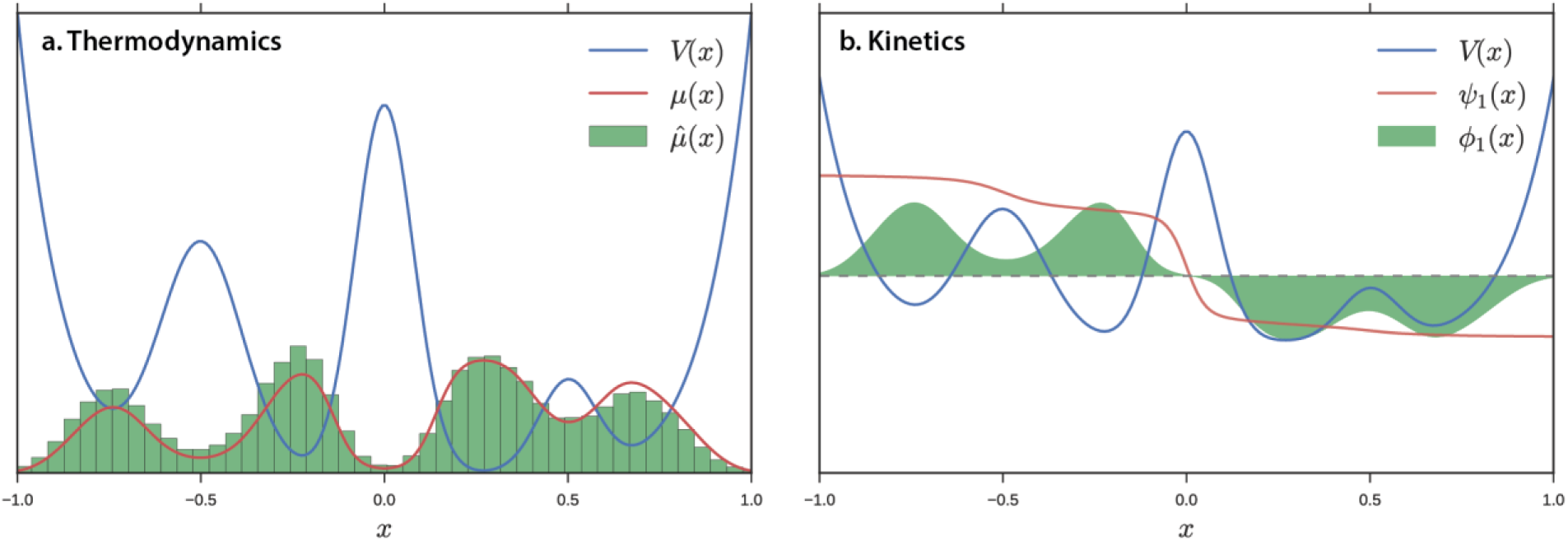
We can characterize a biophysical system by (a) its thermodynamics and (b) its kinetics. Thermodynamic properties can be estimated without regards to time. Here, we estimate the equilibrium distribution by computing a histogram. We model kinetics by estimating the eigenfunctions of the transfer operator (psi) or μ-weighted propagator (phi). Here we plot the slowest dynamical eigenfunction, which represents moving from the left two basins to the right two basins. For this 1D potential energy function, the analytic eigenfunctions are tractable. In analogy to the histogram in (a), we seek a numerical method to evaluate psi or phi from finite data.

tICA is an alternative method for modeling kinetics. Whereas MSMs use indicator functions (bins) to estimate *ψ*, tICA uses linear functions. This corresponds to using hyperplanes (for our 1D potential, a line) to estimate *ψ*, see Figure 2d. Typically for biomolecules, one extracts vector features such as dihedral angles or contact distances to serve as inputs to tICA. Although no longer subject to the limitations of discrete bins (see above), Schwantes and Pande ^11^ noted that the linearity of tICA harshly constrains the solutions. To introduce non-linearity, the authors borrowed a technique from machine learning and introduced a kernel trick. By re-writing the tICA problem only in terms of inner products, researchers can solve the tICA problem in an arbitrarily large, expanded space relative to the original representation *without* explicitly transforming the original representation into this space. By using an appropriate kernel function such as a Gaussian kernel (a type of radial basis function (RBF)), the implicit representation is infinitely large, containing every power of input coordinates (per Taylor expansion of the exponential function). The authors showed that this method could capture the important, slow degrees of freedom from a simulation using many fewer, non-linear coordinates. There are drawbacks to kernel tICA that have precluded it from wide adoption: (1) it is sensitive to hyperparameters. (2) It scales very badly with amount of data. Presciently, Schwantes and Pande ^11^ noted that developments in the kernel learning community could be applied here. In that spirit, we introduce a variant of the Nyström approximation to address the problems identified above (summarized in Table 1). We also show that this novel method connects MSMs and tICA in a novel way.

**Figure 2:**
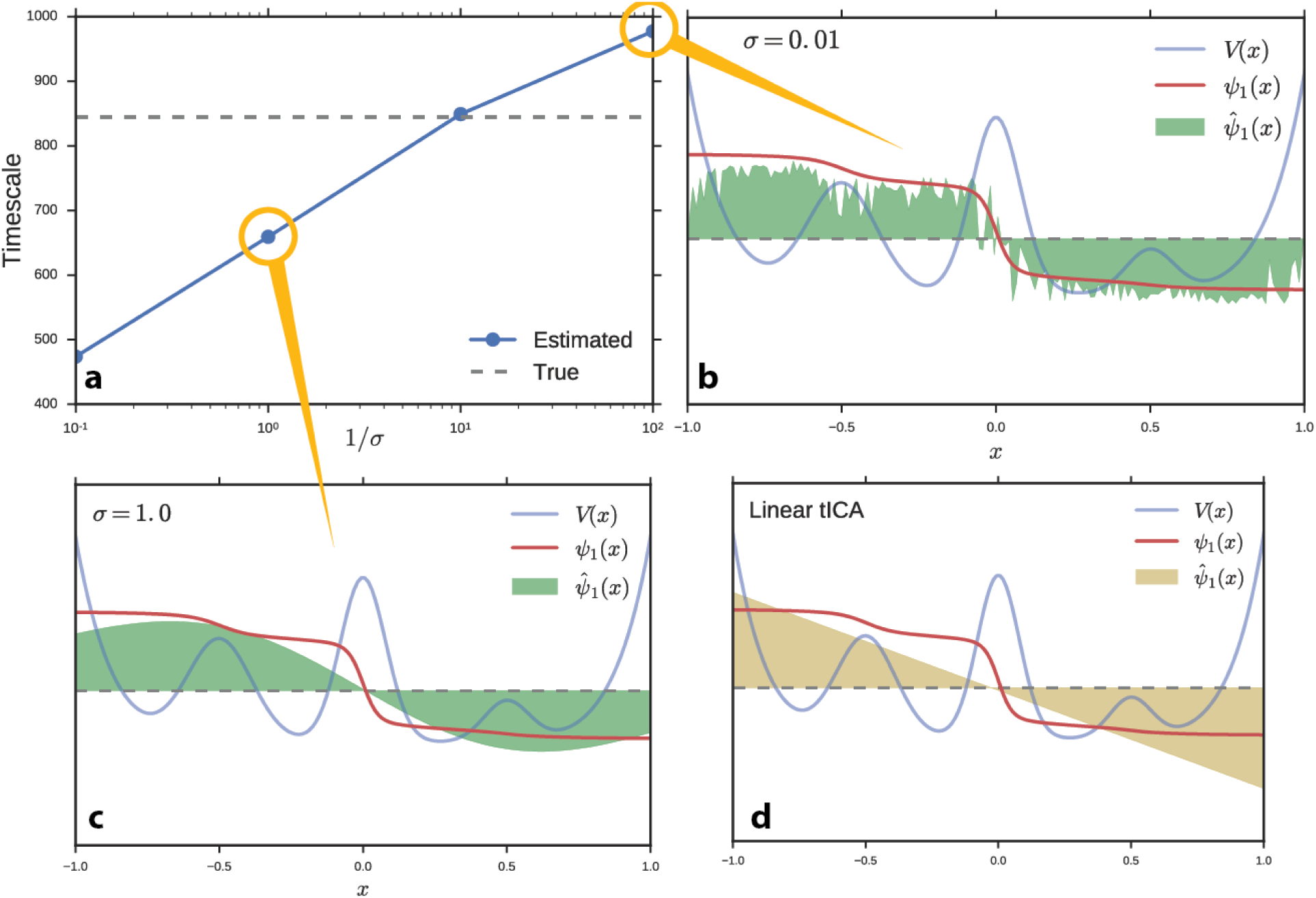
The kernel tICA approach is highly sensitive to the choice of σ (basis function width). (a) The model timescales increase as σ decreases. Note that the variational bound on the model's slowest timescale is easily exceeded as σ shrinks. (b-d) The models numerically estimate (shaded region) the slowest dynamical eigenfunction of the system. (b) A small σ ( width) will overfit to noise. (c) A large width misses nuance in the data and begins to resemble ordinary linear tICA. (d) Ordinary linear tICA.

**Table 1:** Problems with existing solutions motivating the present work

|  | MSM | tICA | Kernel<br>tICA |
| --- | --- | --- | --- |
| Non-linear | Yes | No | Yes |
| Smooth | No | Yes | Yes |
| Tractable | Yes | Yes | No |

## 2 Method

The Nyström method of approximation can be used to speed kernel-trick computations ^12,13^. Instead of computing the Gram matrix *K* ∈ ℝ^mxn^ (where n is the number of data points) in full, we approximate it by 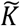 = *K_n,m_*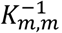*K_m,n_* where *K_n,m_* is constructed by randomly selecting *m* columns of the original matrix. Williams and Seeger ^12^ showed that we can choose *m «η* in practice. Since the limiting step is diagonalizing the Gram matrix, the computational complexity can be reduced considerably from *0*(*n*^3^) in amount of data to *0*(*m^2^n*) ≈ *0*(*n*). We improve upon this approximation by selecting columns (corresponding to data points) of the original Gram matrix according to the result of a (perhaps rough) clustering of the data. This comes at the expense of slightly additional computational cost, but improved coverage of the observed data and added interpretability (see section 2.2). We name the *m* data points corresponding to the selected columns “landmark points”. In practice, we transform the input time-series data by explicitly computing the kernel function to each landmark point and then apply the ordinary linear tICA algorithm in this new space. In this way, the kernelized distance to the landmark points can be thought of as a set of features not dissimilar to dihedral angles or contact distances.

### 2.1 Kernel Function and Distance Metric

Any suitable kernel function can be used with this formalism. In this work, we have constructed our examples with the Gaussian RBF kernel, which is popular in the machine learning community due to (1) its infinite Taylor expansion, giving rise to an infinite dimensional latent feature space and (2) its interpretation as a similarity measure with range [0,1]. It admits one hyperparameter: the basis function width, σ.

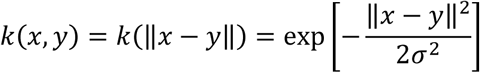

For simple problems in an explicit Euclidean vector space, like the toy potential in section 3.1, the choice of distance metric is simple. A ℓ^2^ norm will suffice. In protein simulations, however, Euclidean norm in the raw Cartesian coordinate space is rarely the best metric of distance. Instead, we require a distance metric that (1) respects translational and rotational degrees of freedom and (2) ignores highly varying but functionally irrelevant conformational changes like hydrogen atom vibrations and solvent motion. Popular distance metrics for MSM construction typically involve transforming the protein Cartesian coordinates into a set of internal coordinates like backbone dihedral angles or distance pairs, and then using the Euclidean norm in this space.

Alternatively, the root mean squared deviation (RMSD) distance metric is a very natural way of thinking about protein conformations. It has been used since the inception of atomistic simulation to describe the difference between conformations (roughly) as the average difference between atomistic coordinates. It takes into account the rotational and translational symmetry of conformations by centering and conformations and reporting the minimum value over rotation. It is a protein-agnostic algorithm that can be applied to any system under study, in contrast with particular distance pairs or dihedral angles which vary in importance between systems. As a disadvantage: RMSD is *only* a distance metric and does not embed a Euclidean vector space. It is impossible to use off-the-shelf machine learning techniques like KMeans clustering or principal component analysis (PCA) or specialized techniques like linear tICA. For this reason, much of the algorithmic advances in MSM construction has focused on vector space features. Landmark kernel tICA does *not* require vector space features (although it works naturally with them as well). We can once again use RMSD in our analysis pipeline.

RMSD is a measure of local distance. With small values (< 3Å), the researcher can be confident that the two conformations are similar. As RMSD values become large (>10Å) the researcher can say that the conformations are different, but the degree of difference is not sensitive to RMSD changes and the ways in which conformations are different cannot be deduced. This gives rise to a rule of thumb for landmark kernel tICA. *σ* can be chosen to be around the cutoff of RMSD’s utility (∼3Å). Other bandwidth selection algorithms from histogram or KDE construction may be applicable here.

### 2.2 Connection to MSMs with soft states

The kernelized distances to landmark points vary between 0 (entirely dissimilar to landmark point) and 1 (equal to the landmark point). We introduce the notion of MSM “states” defined by a centroid conformation and interpret the kernelized distances as partial occupancies in those states. In practice, MSMs are constructed by filling in a “counts matrix” of transitions between labeled states. This is equivalent to computing the time-lagged correlation matrix of state-occupancy vectors of the form | ***k***〉 {0,…, 0,1,0,…, 0} zero everywhere except at the kth position. Define ***K***^(^*^t^*) to be the set of these vectors over time starting at time *t*. Then

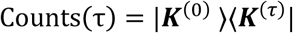

Now take |***k'***〉 to be soft occupancies between zero and one (e.g. the result of kernel featurization). With sufficiently separated landmark points, the vector will take on a form |***k'***〉 = {…,0.1,0.9, 0.1,…}, i.e. all entries close to zero except at the kth position, where it is close to one. The analogous computation:

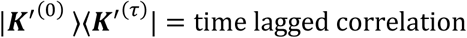
gives the time-lagged correlation matrix, which—when properly normalized by the covariance—gives the tICA eigenvalue problem. This mathematical relation lets us view the landmark kernel tICA model as a Markov state model with “soft” states less susceptible to shot noise and poorly positioned states.

### 2.3 Drawbacks

The tICA components (i.e. eigenvectors) become harder to interpret in this method. When using molecular features like dihedral angles or atom pair distances, the magnitude of the individual values in a tICA eigenvector represents a “relative importance” of that feature to the chosen dynamical mode. Specific amino acids can be colored or otherwise visualized based on their contribution to a particular tICA component, guiding the researcher towards “interesting” regions of a large biomolecule. In this method, the eigenvector values relate to landmark conformations, which may be more difficult to interpret.

The choice of number of landmark points, *m*, as well as the choice of kernel function and associated kernel parameters adds additional tunable hyperparameters, which is never desirable. We address some of these issues in the following sections.

## 3 Results and Discussion

### 3.1 Model quality on a 1D potential

We performed a numerical experiment to determine the effect of hyperparameters on the full kernel tICA solution (without the Nyström approximation) and on the landmark kernel tICA solution. We simulated one hundred Brownian dynamics trajectories on the potential energy function from Figure 1. We learned a kernel tICA model at several values of *σ* and estimated the slowest timescale of the system, Figure 2a. For this simple problem, we can compute the true value of the timescale analytically (dashed line). Interestingly, the true timescale (which should serve as a variational bound in the infinite data limit) is easily exceeded for particularly small values of *σ*. We plot the estimated propagator eigenfunctions at small (Figure 2b) and large (Figure 2c) values of *σ*. Whereas large basis function widths miss nuance and non-linearity in the data and approach the linear tICA limit (Figure 2d), small values result in overfitting to noise (and incorrectly slow timescales). In fact, kernel tICA is highly susceptible to overfitting because each data point is related to every other data point leading to a large number of parameters to fit.

We have demonstrated that relying on the variational principle in the context of finite data would lead us to an improper choice of sigma. We can control for overfitting by using cross validation and the GMRQ score ^10^. By splitting our data into a separate train and test set, we can evaluate how well a model built on the training set can capture the slow dynamics of the test set. We performed this analysis over a variety of choices of *σ* for kernel tICA and landmark kernel tICA, see Figure 3. Note that the landmark approximation shows a marked increase in maximum model quality (as measured by the GMRQ score). It displays a high score over a large range of hyperparameter values, including small values of *σ* where full kernel tICA does especially poorly (c.f. Figure 2b). For large values of *σ* (>10^0^) the full solution performs slightly better. We remind the reader that in this limit, the kernel tICA solution loses its non-linearity and reverts to linear tICA (c.f. Figure 2 c and d). The landmark approximation inherently regularizes the model. By reducing the number of parameters (i.e. number of landmarks), we remove the ability of the model to overfit to noise and spurious data points. In addition to the large computational savings, the landmark ktICA approach also produces better models.

**Figure 3:**
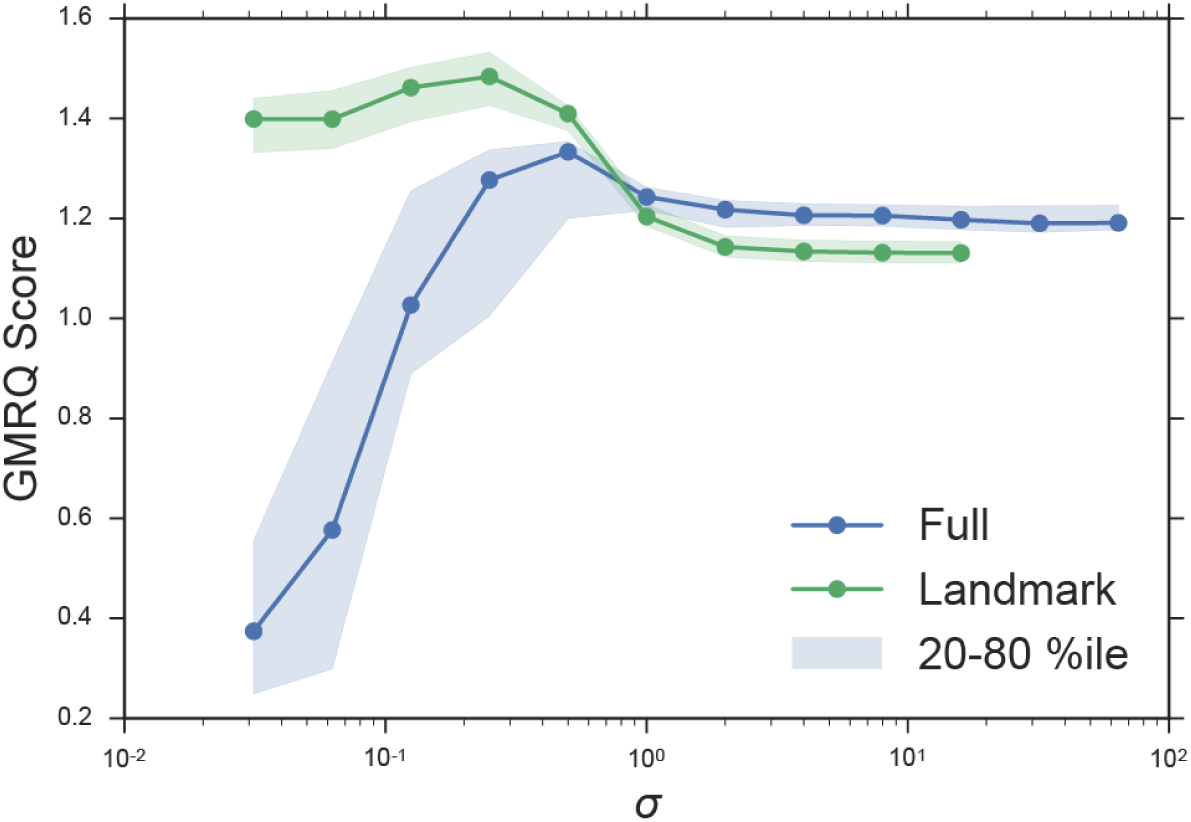
Landmark kernel tICA performs better than full kernel tICA over a large range of σ values. By employing landmarks, we inherently regularize the solution. Full kernel tICA essentially treats every point as a landmark point, which can give rise to solutions like Figure 2b. Additionally, full kernel tICA is computationally unfeasible with O(N^3^) dependence on the number of data points.

Unfortunately, the landmark approximation introduces a new hyperparameter *m*—the number of landmark points. From the perspective that the explicit kernel evaluations are features similar to dihedral angles or contact distances (see Section 2), the MD practitioner can reason that the degree of approximation *m* controls the resolution of the representation of the data and might be chosen to be of the same order of magnitude as the number of dihedrals or number of atoms. Moreover, it suggests that full kernel tICA (*m* = *n*) uses far too many landmark points for a typical MD dataset. As an example, consider a 100-residue peptide simulated for 1ms with frames saved every 1ns. We might expect to describe this system with 10^2^ or 10^3^ features, whereas full kernel tICA would use 10^6^. More stringently, one can use GMRQ cross validation to determine the best selection for this hyperparameter. We performed GMRQ cross validation on the four-well potential dataset across a range of values for *m* and found a weak dependence of score on number of landmark points with a maximum around *m* = 8. This simple toy model does not require a large number of landmark points. The consistently high score across a range of values suggests heuristics may be sufficient for this parameter. In Figure 3, we fixed *m* = 20. In Figure 4, we fixed *σ* = 0.25. Cross validation was 10-fold shuffle-split among 100 trajectories. The two parameters could be simultaneously optimized on a two-dimensional grid search with added computational cost.

**Figure 4:**
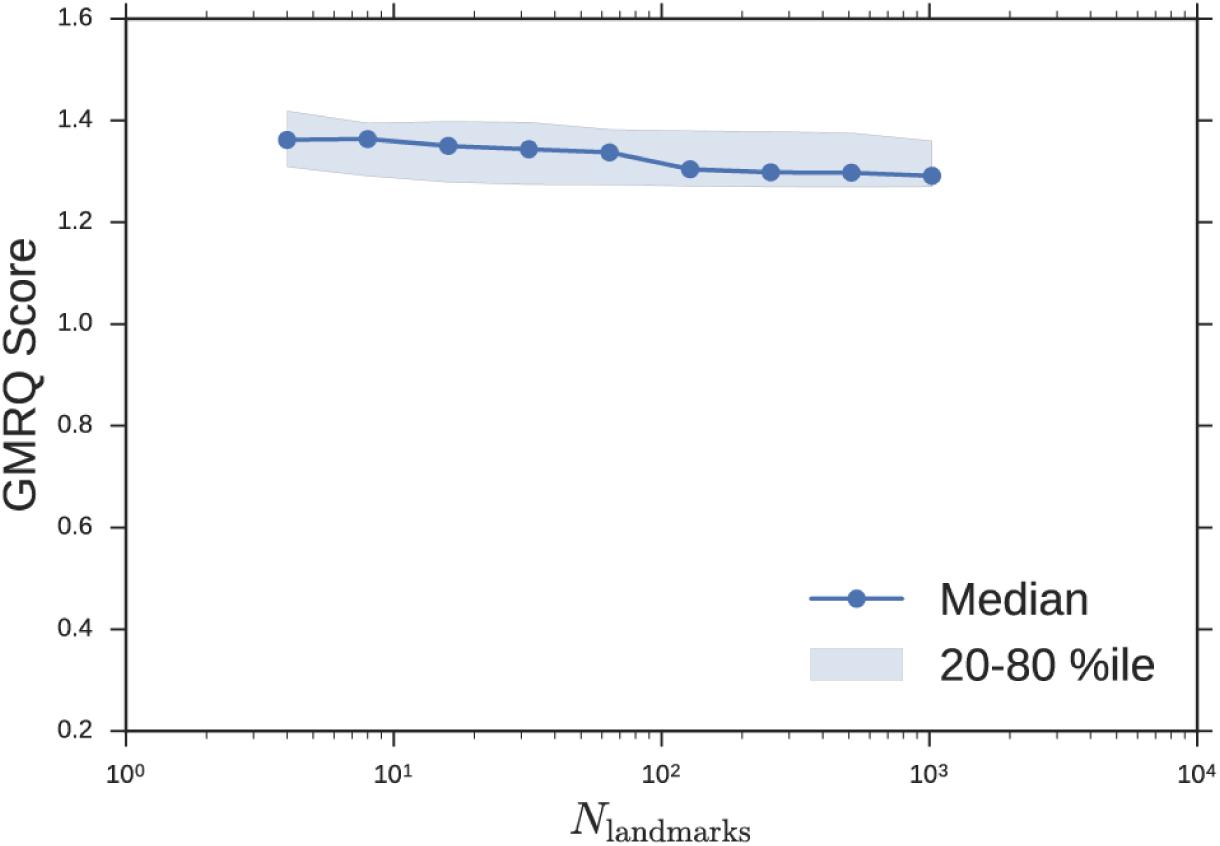
The quality of the model is weakly dependent on number of landmarks, m, for this simple system

### 3.2 A folding coordinate for a small peptide

tICA has been used successfully as an intermediate step during MSM construction. By transforming the data into kinetically-oriented coordinates, defining the indicator-function basis functions via clustering of the data becomes more robust. Building an MSM is still necessary with linear tICA, because the linearity overly constrains the solutions ^6^. Kernel tICA transforms the data into coordinates that can comprise nonlinear movements, and would therefore serve as a better intermediate step for MSM construction. However, by introducing non-linearity to the solutions, we can construct a kernel tICA model that is both accurate and interpretable without building an MSM.

In this example, we re-analyze the fip35 ww domain peptide folding simulations of Shaw et. al.^14^ with landmark kernel tICA. We modeled 200 μs of trajectory data using 500 landmark conformations selected by all-atom RMSD mini-batch kmedoids clustering implemented in MSMBuilder ^15^. Kernel features were computing using the Gaussian kernel with σ = 3Å. The tICA model was computed with a lag-time of 10 steps or 1ns. The large spectral gap in tICA timescales (Figure 5, right) confirms that the slowest timescale is accurately captured by only one landmark kernel tIC. Although this is a loaded term ^16^, we declare this tIC to be the folding reaction coordinate. We can negative-log-histogram the data along this coordinate to estimate the energy profile for folding, Figure 5. The energy landscape shows a two-state behavior. The global minimum is the folded structure (Figure 5, inset, left). The metastable basin is a collection of unfolded conformations; we show the unphysical structural mean of a number of these conformations (Figure 5, inset, right).

**Figure 5:**
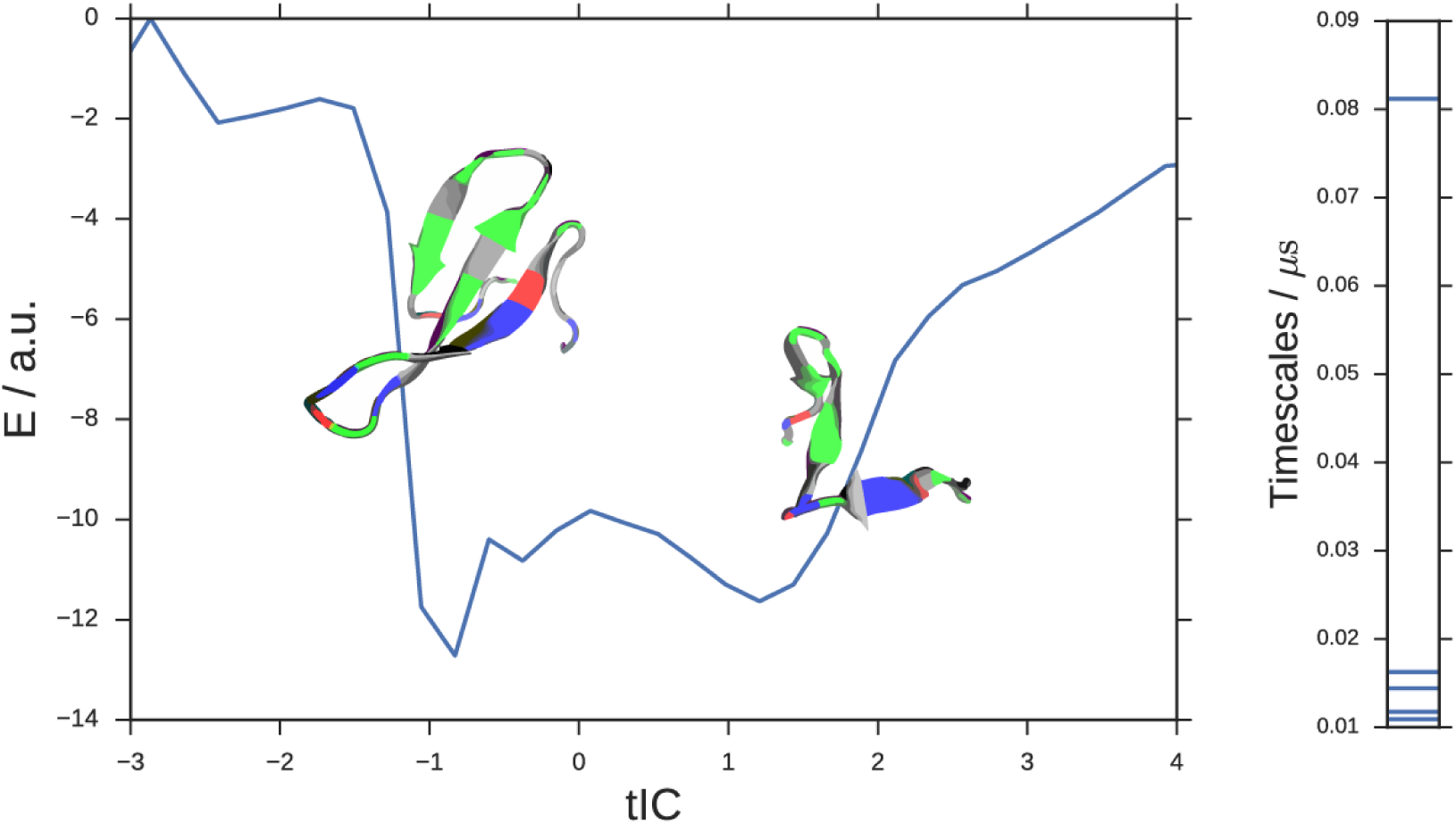
An energy landscape for the fip35 ww peptide. The analysis captures the minimum-energy folded state and an unfolded basin without hand-picking any coordinates or reference structures. Example conformations are overlaid on their tIC value. The learned kernel tICA model shows a large spectral gap (model timescales, right) which supports the projection of the dataset onto a single tIC coordinate.

We can generate a trajectory (movie) by selecting points along the tIC coordinate and drawing conformations from the raw data. See the SI for the fip35 kernel tIC folding trajectory. Note that these trajectories are often unphysical due to the naïve way we draw conformations along the tIC. In particular, the large structural changes that are all consistent with the entropically-dominated unfolded basin are often selected in adjacent time points. In the movie, we have smoothed the trajectory; in the unfolded region, this often results in atoms being averaged on top of each other. Further work can be done to produce more realistic and visually appealing movies, perhaps by selecting conformations consistent with the desired tIC value *and* similar to the previous frame.

### 3.3 An activation coordinate for conformational change

We apply the new method to a potassium ion channel that was previously shown to undergo a large conformational change between a compressed, down state and a stretched, up state. Similar to the peptide example, we recaptured the up-down dynamics with a single landmark kernel tIC. We used 400 μs of aggregate Folding at Home trajectory data to extract 500 alpha-carbon RMSD landmark points. Once again, we used the Gaussian kernel with σ = 3Å. The tICA model was computed at a lag time of 10 steps or 9.6 ns. In Figure 6, we histogram the data along the learned coordinate to estimate a free-energy diagram for the up-down transition. The right basin is the stretched, up state with intracellular helices splayed outwards. The left well comprises the compressed, down state with rigid helices. For this system, there is an additional slow process within the compressed basin, identifying a metastable down-like state noted in the original study. See figure S1 for a detailed comparison of the two down-like states. By sampling representative conformations along this coordinate, we prepared a trajectory showing the up-down process, see the movie in the SI.

**Figure 6:**
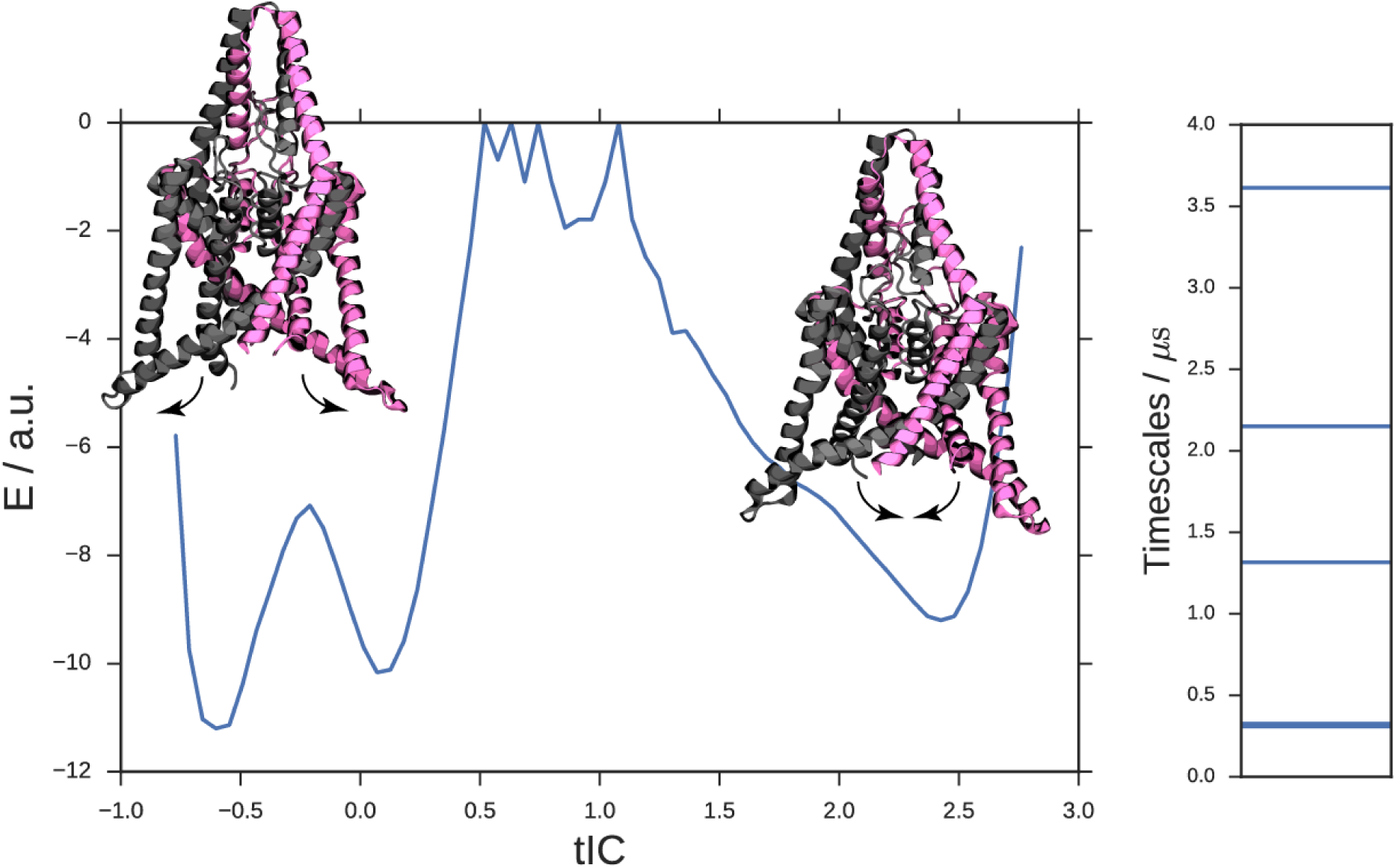
Landmark ktICA learns the up-down coordinate from a simulation of the TREK-2 leak channel. The data is clearly separated into two large wells. The right well represents the “up” state resulting from membrane stretch. The left well represents the down, compressed state. The down contains an intrastate barrier separating what was previously identified as a down-like intermediate. See figure S1 for a detailed comparison of the two down basins.

## 4 Conclusions

Landmark kernel tICA is the culmination of years of research into using MSMs, tICA, and the transfer operator formalism for understanding conformational dynamics. The first key insight was when Swope and Pitera ^4^ applied the theory of Markov chains to protein dynamics. As researchers applied this formalism, it became clear that state-space definition was crucial for constructing accurate MSMs. By applying the theory behind principal component analysis (PCA), several groups identified tICA as a useful dimensionality reduction for defining states in MSM construction. With the revelation that tICA and MSMs differ only by choice of basis, Schwantes and Pande ^11^ extended tICA using the kernel trick to build models of protein dynamics directly. We revisit MSMs by re-introducing the notion of “states”, this time with fractional occupancy based on kernel distance to the state centroid. In so doing, we introduced landmark kernel tICA which can be considered simultaneously (1) a regularized, computationally-tractable nonlinear tICA model and (2) a Markov state model with “soft” states. This “best-of-both-worlds” approach yields highly accurate models that can be interpreted as MSMs *or* via projection onto reaction coordinates. We introduce this method with an eye towards improving existing methodologies that rely on robust projection of data. In particular, accelerated sampling schemes like umbrella sampling or metadynamics make strong assumptions about orthogonal degrees of freedom: the chosen reaction coordinates over which we accelerate are “slow” but every degree of freedom orthogonal to them must equilibrate must faster (e.g. within an umbrella window timeframe). Without increasing the number of accelerated coordinates (which typically scale poorly), we anticipate that the present work can be used to choose coordinates for which this assumption is more likely to be true. Due to improved model quality, high interpretability, and computational tractability, the landmark kernel tICA approach can be widely applied to the analysis of dynamical biomolecules.

## 5 Acknowledgements

We thank Christian Schwantes, Robert McGibbon, Muneeb Sultan, and Carlos Hernandez for inspiration and helpful discussions. We thank Keri McKiernan, Muneeb Sultan, and Brooke Husic for critical feedback on the manuscript.

We thank the National Institutes of Health grant number NIH R01-GM62868 for funding.

We graciously acknowledge D. E. Shaw Research for providing access to the fip35 folding trajectory datasets.

## 5.2 Author contributions

MPH designed and performed the research and wrote the manuscript. VSP supervised the research and edited the manuscript.

## Supporting Information

**Figure S1:**
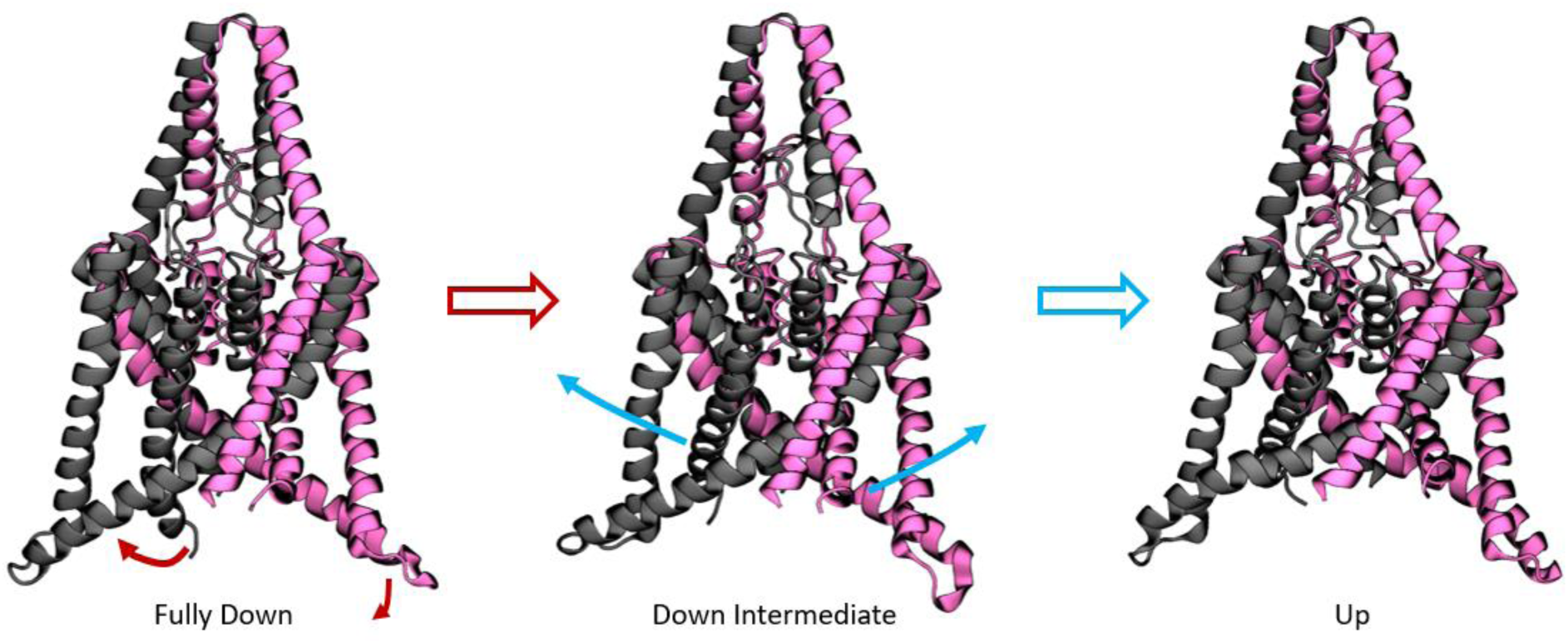
Structural details between all three wells identified in Figure 6. The fully down state (left) has all helices in their down-most configuration. In the down-intermediate state (middle), one of the inner pore helices (grey) has moved up and there is local unfolding on the outer helix. In the up state (right), all helices are in their up-most configuration.

